# The impact of the “World’s 25 Most Endangered Primates” list on scientific publications and media

**DOI:** 10.1101/541706

**Authors:** Daphné Kerhoas, Alberto Acerbi, Amanda D. Webber, Gráinne McCabe, Russell A. Mittermeier, Christoph Schwitzer

**Affiliations:** Bristol Zoological Society, Bristol, United Kingdom; School of Innovation Science, Eindhoven University of Technology, The Netherlands; Global Wildlife Conservation, Austin, Texas, USA

**Keywords:** Primate conservation, conservation outreach, bibliometric analysis, digital media, social media

## Abstract

Assessing the impact of conservation campaigns is of high importance for optimizing the use of limited resources. Lists of threatened species are often used as media outreach tools, but their usefulness is rarely tested. We investigated whether the inclusion of a species in the list “World’s 25 Most Endangered Primates”, published biannually by the International Primatological Society, the International Union for Conservation of Nature’s Species Survival Commission Primate Specialist Group, and Conservation International from 2000, had an effect both on scientific publications and on the general public. We analyzed a database of 40 million articles from major scientific publishers (Elsevier, Springer, Nature, Plos, Pubmed, Biomed Central) finding an increase in the number of papers mentioning a species after its inclusion in the list. We also analyzed media penetration (data from Google News), and online interest (data from Google Blogs, Twitter, and Google Trends), collecting daily data for one month before and one after the official launch of the 2014-2016 list (24^th^ November 2015). The results show a short spike of interest on Google News and Twitter but no long term effect, indicating a limited effect on the general public. Our results are important for the understanding of the impact of current conservation campaigns and to provide strategies for future campaigns.

## 1 Introduction

It is widely recognised that the internet could be a useful tool to understand and explore public interest around a specific event or general issues. Large volumes of data are freely and easily accessible providing a cost effective way of analysing trends and attitudes across a broad spectrum of the public (see Proulx et al. 2014, Anderegg & Goldsmith 2014, Cha & Stow 2015, Soriano-Redondo et al. 2017). For example, the developing field of culturomics examines large online databases of word frequencies using offsite metrics that can then be used to understand or predict behaviour or processes. One of the best-known examples is Google Flu Trends which utilises internet search data to track and ultimately plan responses to flu outbreaks (Dugas et al. 2013). This relies on the google search engine, the world’s most commonly used search engine with 80% of the global market share (Netmarketshare 2017). Whilst it has been argued that these online tools may have less bias than traditional methods (Soriano-Redondo et al. 2017) and are particularly effective if triangulated with other tools (Proulx et al. 2014), it should be noted that models need to be adapted. Despite historical accuracy, in 2013, Google Flu Trends did not accurately predict peak levels of flu in the US (Butler 2013).

Despite the growing use of digital resources in other areas, bilbiometrics, social media, and internet search data are still little used in conservation research (Proulx et al. 2014, Cha & Stow 2015). A small number of studies have, however, used online sources to examine trends in public interest in environmental issues (Ficetola 2013, Mccallum & Bury 2013, Soriano-Redondo et al. 2017), and monitor ecosystem services and trade (Galaz et al. 2010, Ladle et al. 2016). Proulx et al. (2014), for example, tracked biological processes and distribution, e.g. pollen and spread of invasive species, and the relationship with public interest. Furthermore, online tools have been used to measure public interest (Nekaris et al. 2013) and potential changes in opinion following key media events including ‘climate gate’ and the death of Cecil the Lion (Anderegg & Goldsmith 2014, Cha & Stow 2015, Carpenter & Konisky 2017). The potential for digital data to assist with understanding support, or a lack thereof, for conservation initiatives has not been yet fully explored (Ladle et al. 2016, Soriano-Redondo et al. 2017)

Since 2000, the International Union for Conservation of Nature’s Species Survival Commission (IUCN SSC) Primate Specialist Group, the International Primatological Society, and Conservation International have biennially published the “World’s 25 Most Endangered Primates” (also known as “Top 25 list” or “Primates in Peril”; hereinafter referred to as “Top 25”). This report highlights twenty-five of the most threatened primate species with the aim of attracting attention and action from the scientific community, relevant governments, and the public. As such, inclusion in the list is not based on the actual conservation status of the primate species, but most are also officially classified as “threatened”. The list is produced by the world’s leading primatologists and field researchers who have first-hand knowledge of the ongoing evolution of threats to primate species; more than 250 experts have been involved in compiling the last five iterations of the publication. The number of species included in this list is evenly distributed between 4 geographical regions (Neotropics, Africa, Madagascar and Asia). Whilst the potential to increase scientific interest and raise the profile of these animals is clear, the actual impact of the Top 25 has never been tested.

The aim of this research is to evaluate the scientific output and media penetration of the Top 25 list. We investigated whether the inclusion of a species in the list had an influence on the number of peer-reviewed articles published on that species in the following years. This is of vital importance as policy-makers and funding agencies rely mostly on scientific reports. We also examined whether the list was an effective communication tool for conservation by analysing media output following the publication of the Top 25 in 2015.

## 2 Methods

### 2.1 Scientific publications

We tested the impact of the mention of a species on the Top 25 list on scientific publications (see Table A1 in the Online Appendix for all species included, and the year of their mentions). We have included in this analysis a total of 37 species that were mentioned at least once in the Top 25 list from 2000-2002 to 2010-2012 (6 lists overall of 25 primate species each). We excluded species that were mentioned in the lists of 2012-2014 and 2014-2016 (as there is not enough post-mention data to assess the impact). Each species was considered separately and included once in the analysis.

We used 74 control primate species (see Table A2 in the Online Appendix) that have never been mentioned in any of the Top 25 lists released to account for a possible bias of an overall increase of publications through time. These control species were chosen randomly, with the constraint of being evenly distributed in the 4 biogeographical regions (Africa, Asia, Neotropics and Madagascar).

We extracted data from 40 million articles published from 1994 to 2014 in six major scientific publishers (PLOS, BMC, Elsevier, Springer, Nature and Highwire/Pubmed; see Table 1). The data were extracted from the publisher databases using custom-written python interfaces to the API they provided. We extracted all articles in which the Latin name of a species that was either included in the Top 25 list (n=37 species) or of control species (n=74 species). We used the Latin name for both Top 25 species and control species as the common name may have changed over the years and scientific articles always list the Latin name when a species is first mentioned. Data from the archives of these publishers were extracted in February and March 2014.

**Table 1:** List of publishers used for the data mining analysis on scientific publication. Search of the species name (either Top 25 species or control) was done either on the full text or on the keywords of scientific articles.

| Publishers name | Search type | Total articles searched | Top 25 species match | Control species match |
| --- | --- | --- | --- | --- |
| PLOS | Full text | 53,500 | 213 | 148 |
| BMC | Full text | 189,955 | 149 | 132 |
| Elsevier | Full text | 11,000,000 | 4,265 | 6,805 |
| Springer | Keywords | 5,000,000 | 66 | 36 |
| Nature | Full text | 500,000 | 211 | 259 |
| HighWire/PubMed | Full text | 23,000,000 | 2,565 | 6,276 |
| **Total** |  | **39,743,455** | **7,469** | **13,656** |

We used a Bayesian structural time-series model that estimates the causal effect of a designed intervention on a time series, given a baseline model of the expected trend (Brodersen et al. 2015) in R software (R Core Team 2014). For each species (Top 25 and control) we compiled a count of the number of scientific articles per year from 1994 to 2014. For species mentioned more than one time in the Top 25, the intervention tested is the period of time from the first to the last mention in the list. We used the average number of scientific publications of the control species trend as baseline. We also ran the same analysis using only control species that were classified as “threatened” (IUCN, 2017) as a control baseline (37 out of 74). This allows us to account for the conservation status of control species which may influence the number of publications.

One key assumptions of this analysis is that the set of control time series should be predictive of the outcome time series in the pre-intervention period. In our case, it is fair to assume that a general rise of publication as observed for control species is to be predicted for the species of the Top 25 before their mention in the list. A second assumption is that the control time series must not have been affected by the intervention (Brodersen et al. 2015). It is unlikely that the scientific publication on a control species, never included in a Top 25 list, would be affected by the release of a biennial Top 25 list.

### 2.2 Media penetration

The Top 25 list for 2014-2016 was decided on the 13^th^ of August 2014 and officially released on the 24^th^ November 2015. We tracked, starting approximately one month before the day of the official launch and for one month after (21/10/15 to the 28/12/15), the presence of a series of keywords (the title of the list itself and related keywords, e.g. “endangered primates”, “primates in peril”, “Top 25 primates”) and the scientific and common names of the 25 primate species included in the list, (e.g. Sumatran orangutans, *Pongo abelii* and red ruffed lemur, *Varecia rubra*, cf. Table A3 in the Online Appendix) on a daily basis. The two data (title/keywords and species names) are considered separately in the analysis. We assessed the penetration of the Top 25 in traditional media (tracked through Google News), social media (through Twitter), blogs (through Google Blogs Search), and the interest of the general public (tracked through Google Trends). Google News is a free news aggregator that selects syndicated web content such as online newspapers in one location for easy viewing. Twitter is a social network where users post messages that can be read by an unregistered person and it has more than 319 million monthly active users as of 2016. Google Blog Search is a service to search blogs content with an identical process to Google Search. Google Trends, that provides data on individual searches in Google, shows how often a term is searched for relative to the total number of searches worldwide.

As in the previous analysis, we used a Bayesian time series analysis (Brodersen et al. 2015). In this analysis we did not consider any control species given that we did not expect any general increasing trend as we did for the scientific publications. We ran the analysis for a post intervention period both of one week and one month, in order to examine the duration of the possible effect.

The data used in the analysis are available in an Open Science Framework repository at https://osf.io/e7ymv/s

## 3 Results

### 3.1 Scientific publications

We found 4,545 scientific articles that contained at least once the Latin name of the 37 primate species that were included in one of the six Top 25 lists from 2000-2002 to 2010-2012. In addition, 13,656 scientific articles contained at least once the Latin name of the 74 primate control species.

Twenty two out of 37 species (59%) had an increase in scientific publications following their inclusion in the Top 25 list (Figure 1). For 11 species there was no identified effect, and 4 species had a decrease in publications following inclusion in the Top 25 list. The four species with the most positive impact were the mountain gorilla (*Gorilla beringei beringei*), the drill (*Mandrillus leucophaeus*), the golden lion tamarin (*Leontop-ithecus rosalia*) and the black snub-nosed monkey (*Rhinopithecus bieti*). The four species that suffered a decline in publication were the brown spider monkey (*Ateles hybridus brunneus*), the Miller’s langur (*Presbytis hosei canicrus*), Miss Waldron’s red colobus (*Procolobus badius waldroni*) and the north-west Bornean orangutan (*Pongo pygmaeus pygmaeus*). There were no significant differences between species mentioned once (n=21) or several times (n=16) in the Top 25 list (two-tailed Mann-Whitney U-test, U=173, p=0.8916; Figure A1 the in Online Appendix).

**Figure 1:**
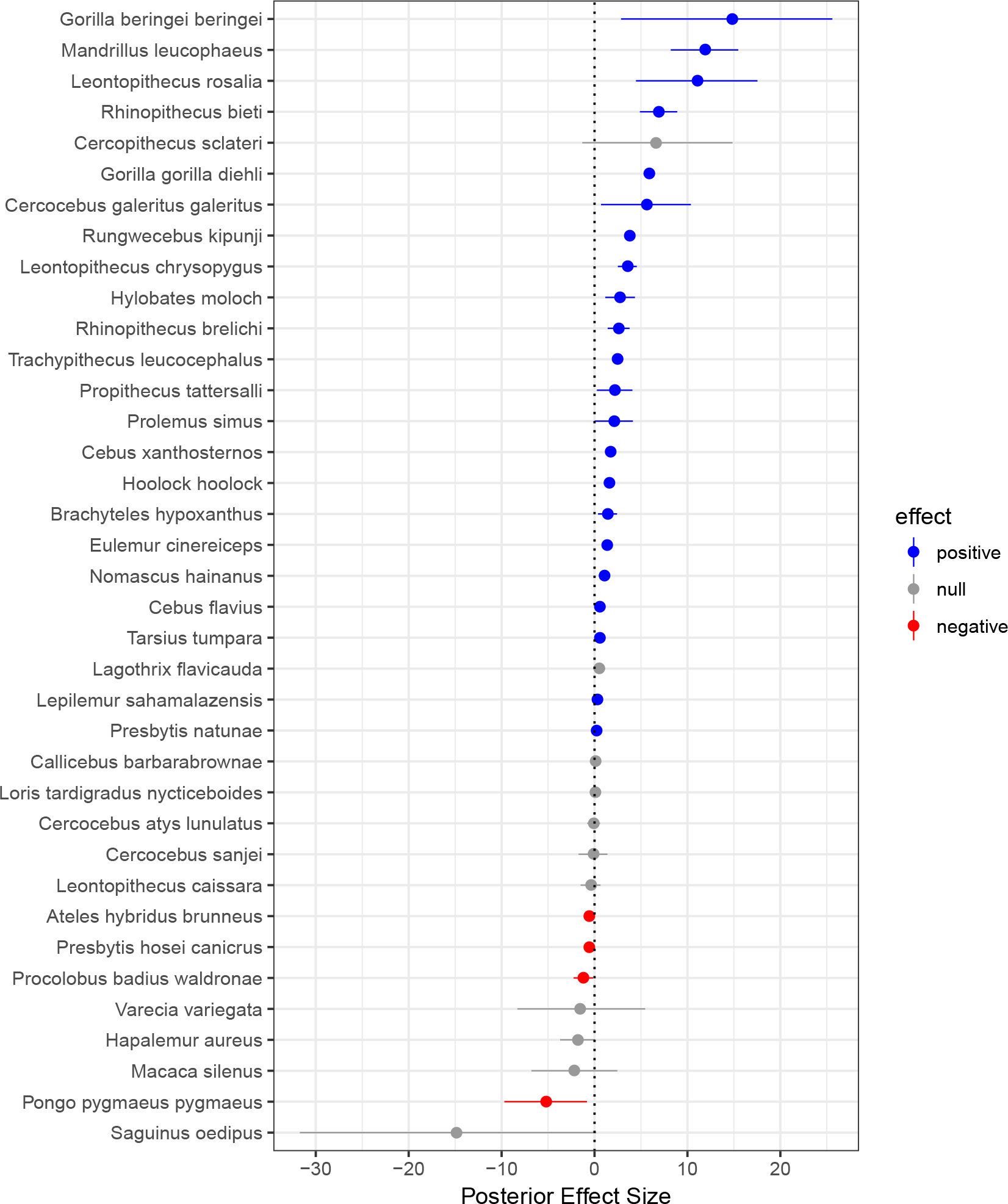
Effect of Top 25 inclusion on scientific publications. Posterior effect size of Causal Impact analysis for each Top 25 primate species included in the 6 Top 25 lists from 2000-2002 to 2010-2012 on scientific publications containing at least once their Latin names. Effect size containing only positive values are in blue, containing both positive and negative value are in grey and containing only negative value are in red.

When using only the control species that were classified as “threatened” (IUCN, 2017) as a baseline to control for publication bias the results were even stronger, with 25 species out of 37 (67.6%) demonstrating an increase in publication rates following their inclusion in the Top 25 list (Figure A2 in the Online Appendix). Twelve species were not affected by their mention in the list and none suffered a decrease in presences in scientific publications after inclusion on the Top 25 list.

### 3.2 Media penetration

#### 3.2.1 Google News

During the pre-intervention period, we collected a total of 296 mentions of the Latin name of the species included in the Top 25 list and 27 mentions of the title/keywords. During the post-intervention period, Latin name of species in the Top 25 list were mentioned 427 times and the keywords 161 times.

When considering a post period of one week, we found a net significant increase of mentions of the common or Latin name of species included in the 2012-2014 Top 25 Most Endangered Primate list (Table 2). However, with a post-intervention period of one month, although the intervention appears to have caused a positive effect, this effect is not statistically significant (Figure 2).

**Table 2:** Latin and Common species names in media. Causal impact analysis results for search of Latin and Common species included in the Top 25 list 2012-2014 on Google News, Google Blogs and Twitter with a pre-period before the official lunch of one month and a post-intervention period after the official launch of either one month or one week. The absolute average effect is the estimated average causal effect across post-intervention period. The absolute cumulative effect is determined as the difference between the predicted and actual value, i.e., the additional publications following the inclusion in the Top 25 list. The relative effect shows the percentage of increase or decrease following the intervention from the predicted values. All effects are reported with their 95% CI.

| Media type | Post-intervention period | Absolute average effect | Absolute cumulative effect | Relative effect in % |
| --- | --- | --- | --- | --- |
| News | month | 3.5 [-3.5, 11] | 121.5 [-122.6, 393] | 40 [-40, 129] |
|  | week | 36 [24, 48] | 291 [189, 381] | 415 [269, 543] |
| Blogs | month | 1.8 [1.7, 1.9] | 64.0 [61.1, 67.0] | 6342 [6058, 6639] |
|  | week | 7.1 [7, 7.2] | 56.8 [56, 57.8] | 24296 [23834, 24748] |
| Twitter | month | 4 [-3.4, 11] | 141 [-119.8, 399] | 23 [-19, 64] |
|  | week | 17 [3.6, 29] | 133 [28.5, 230] | 93 [20, 160] |

**Figure 2:**
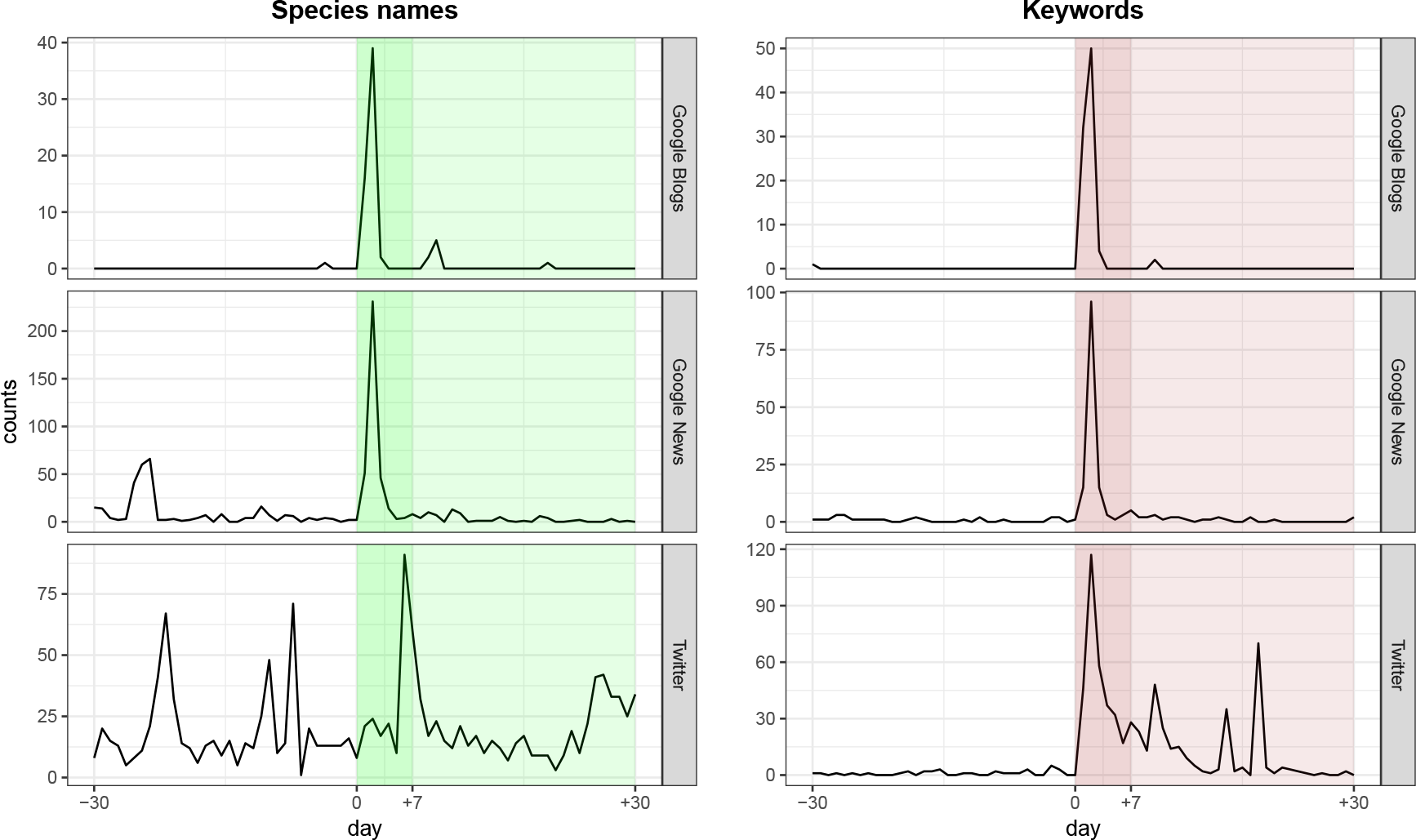
Effect of Top 25 inclusion on media. Counts of mentions on Google Blogs, Google News and Twitter of Latin name species and keywords related to the list one month before and one month after the official launch of the Top 25 list (24th of November 2015). The post-intervention period (following the launch) of one month and of one week are highlighted.

When we considered the keywords associated with the Top 25 list we found that there was a significant effect of the official launch on the use of these keywords in Google News, both considering a post-intervention period of one week and of one month (Table 3).

**Table 3:** Top 25 related keywords in media. Causal impact analysis results for search of keywords (e.g. top 25 primates, primate in peril) included in the Top 25 list 2012-2014 on Google News, Google Blogs and Twitter with a pre-period before the official lunch of one month and a post-intervention period after the official launch of either one month or one week. The absolute average effect is the estimated average causal effect across post-intervention period. The absolute cumulative effect is determined as the difference between the predicted and actual value, i.e., the additional publications following the inclusion in the Top 25 list. The relative effect shows the percentage of increase or decrease following the intervention from the predicted values. All effects are reported with their 95% CI.

| Media type | Post-intervention period | Absolute average effect | Absolute cumulative effect | Relative effect in % |
| --- | --- | --- | --- | --- |
| News | month | 3.8 [3.4, 4.2] | 1133.2 [117.7, 148.2] | 480 [424, 534] |
|  | week | 17 [16, 17] | 134 [128, 139] | 2100 [2015, 2182] |
| Blogs | month | 2.5 [2.4, 2.5] | 86.1 [83.2, 88.9] | 4446 [4295, 4590] |
|  | week | 11 [11, 11] | 86 [84, 87] | 19152 [18901, 19379] |
| Twitter | month | 17 [16, 17] | 588 [568, 610] | 1726 [1666, 1790] |
|  | week | 44 [43, 45] | 350 [343, 358] | 4486 [4394, 4587] |

#### 3.2.2 Google Blogs

The Latin name of the species included in the Top 25 list and keywords relating to the list were both mentioned only once during the pre-intervention period in Google Blogs. During the post-intervention period, Latin name of species in the Top 25 list were mentioned 65 times, and the keywords 88 times.

We found that with both a short and long post-intervention period there was a significant effect of the Top 25 list official launch on the mention of Latin and common names of species (Table 2) on the use keywords (Table 3) included in this list (Figure 2).

#### 3.2.3 Twitter

Latin and common name of species were included in tweets 621 times during the pre-intervention period. Keywords associated with the Top 25 list were sporadically used in comparison, with a total of 33 tweets. For the post-intervention period, there were 768 mentions in tweets including Latin or common names of species included in the Top 25 list and 622 mentions of the Top 25 associated keywords.

Our analysis of the number of tweets and retweets following the Top 25 list launch in 2015 yielded similar results to Google News (Figure 2). When considering the species name there was an effect of the launch on mentions on twitter in the one week-post intervention period, but no effect in the one month period (Table 2). The analyses on keywords yield significant results for both period lengths (Table 3).

#### 3.2.4 Google Trends

After looking up on Google Trends the different species included on the Top 25 2014-2016 (using either the scientific and common names), we find that there was no impact of the Top 25 being released on the number of individual searches in Google. In fact, there were too few individual searches on Google to use this dataset in further analysis. Thus we did not analysed further using the data extracted on Google Trends.

## 4 Discussion

We found that inclusion in the “World’s 25 Most Endangered Primates” list had a positive effect on the number of scientific papers published on the featured primate species. This is encouraging, and it suggests that the use of this type of report can drive scientific interest for these threatened species. Furthermore, as policy-makers and funding agencies rely on scientific reports, this could have a direct positive impact on the conservation of these primates. This result is, in some ways, unsurprising as some of the scientists publishing on these species are going to be those who contribute to the formulation of the Top 25 list. It is difficult to untangle the direction of impact e.g., is inclusion driving publications or is the author’s involvement with the list driving inclusion? The lack of causal inference is a recognized limitation with this type of online data (Proulx et al. 2014, Nghiem et al. 2016) and suggests the need for further research.

The primate species that suffer a decrease of publications following the inclusion are from two distinct regions, namely the Neotropics and Asia. The inclusion on the list of an ape species is found to either improve greatly or decrease scientific publications on these species (respectively the mountain gorilla and the north-west Bornean orangutan).

Examination of media penetration highlighted a significant increase in news articles focusing on species included in the Top 25 list, but this was not sustained for a month after publication of the report. This has also been seen in other studies where there tends to be a short term interest in the issue that is not sustained e.g., the killing of Cecil the lion (Carpenter & Konisky 2017) or media events regarding climate change (Anderegg & Goldsmith 2014). The short spike of interest might be due to high news turnover.

Interestingly, there was a significant increase in attention in Google Blogs for species that had been included in the Top 25 list. This result may mostly be due to the absence of any keywords and species name in the pre-period. Thus, even with a few mentions in any blogs found in Google after the official launch, the analysis may yield a significant effect of the intervention on the data collected. The sustained interest, i.e., after one month, may also be a reflection of the longer timeframe required to extract information from news sites, write and publish blogs. However, it also suggests that direct engagement with key influencers and bloggers would have potential to increase the reach of news regarding key conservation events. We found too few data points on Google Trends to justify an analysis. This may mean that the general reader about the Top 25 online either already know the list or do not research its meaning on Google.

This result was also seen in the social media analysis where there was an increase in attention on included species one week after launch of the Top 25 list. However, we do not know whether there was a long term interest within the social media sphere. In fact, conservationists need to understand how to use social media effectively and engage with their audience (Papworth et al. 2015). Simply releasing reports or updates on to Twitter is not enough for a sustained impact and suggests there is the need to intensify engagement and support with a social media friendly communication tool (such as videos). It also requires the collaboration of conservation partners and scientists to gain any traction within social media. In its current form, the Top 25 list is, therefore, not effective as a communication tool to the public. However, with a more structured, multiple release to a developed online community we believe its impact could be improved. The use of onsite metrics will also be important to understand public interest and improve conservation information penetration (Soriano-Redondo et al. 2017)

In conclusion, use of offsite metrics to examine the impact of a conservation intervention provides an important insight into scientific and public interest. This is necessary to drive future communication in this area (Anderegg & Goldsmith 2014, Nghiem et al. 2016) However, there are limitations of this method which need to be taken into account (Ladle et al. 2016). For example, the reliance on English speaking search engines has the potential to skew the data as there are other online tools used extensively in other countries; whilst Baidu has only a 6% global market share, it has 70% of the market share in China (Statcounter 2017).

The “World’s 25 Most Endangered Primates” publication appears to fulfil its aim on attracting attention and action from the scientific community. It has a positive impact on scientific publications and, by association, research into these threatened species. Impact on governments is harder to ascertain and was not the focus of this study. There seems to be little impact, however, on attracting the attention of the general public and we would suggest that this becomes a focus of the publishing team going forward.

## Acknowledgements

This work has been funded by the Bristol Zoological Society and by the Margot Marsh Biodiversity Foundation. We are grateful to Anna Egerton, Alexia Balatsoukas, and Claire Drury for help in data collection, and to Anthony Rylands for comments to the manuscript.

## References

Anderegg, W. R. & Goldsmith, G. R. (2014), ‘Public interest in climate change over the past decade and the effects of the ‘climategate’ media event’, Environmental Research Letters 9(5), 054005.

Brodersen, K. H., Gallusser, F., Koehler, J., Remy, N., Scott, S. L. et al. (2015), ‘Inferring causal impact using bayesian structural time-series models’, The Annals of Applied Statistics 9(1), 247–274.

Butler, D. (2013), ‘When google got flu wrong’, Nature 494(7436), 155.

Carpenter, S. & Konisky, D. M. (2017), ‘The killing of cecil the lion as an impetus for policy change’, Oryx pp. 1–9.

Cha, Y. & Stow, C. A. (2015), ‘Mining web-based data to assess public response to environmental events’, Environmental pollution 198, 97–99.

Dugas, A. F., Jalalpour, M., Gel, Y., Levin, S., Torcaso, F., Igusa, T. & Rothman, R. E. (2013), ‘Influenza forecasting with google flu trends’, PloS one 8(2), e56176.

Ficetola, G. F. (2013), ‘Is interest toward the environment really declining? the complexity of analysing trends using internet search data’, Biodiversity and conservation 22(12), 2983–2988.

Galaz, V., Crona, B., Daw, T., Bodin, Ö., Nyström, M. & Olsson, P. (2010), ‘Can web crawlers revolutionize ecological monitoring?’, Frontiers in Ecology and the Environment 8(2), 99–104.

Ladle, R. J., Correia, R. A., Do, Y., Joo, G.-J., Malhado, A. C., Proulx, R., Roberge, J.-M. & Jepson, P. (2016), ‘Conservation culturomics’, Frontiers in Ecology and the Environment 14(5), 269–275.

Mccallum, M. L. & Bury, G. W. (2013), ‘Google search patterns suggest declining interest in the environment’, Biodiversity and conservation 22(6-7), 1355–1367.

Nekaris, K. A.-I., Campbell, N., Coggins, T. G., Rode, E. J. & Nijman, V. (2013), ‘Tickled to death: analysing public perceptions of ‘cute’ videos of threatened species (slow lorises–nycticebus spp.) on web 2.0 sites’, PloS one 8(7), e69215.

Netmarketshare (2017), ‘Market share reports: search engines’, available at: www.netmarketshare.com.

Nghiem, L. T., Papworth, S. K., Lim, F. K. & Carrasco, L. R. (2016), ‘Analysis of the capacity of google trends to measure interest in conservation topics and the role of online news’, PloS one 11(3), e0152802.

Papworth, S. K., Nghiem, T., Chimalakonda, D., Posa, M. R. C., Wijedasa, L., Bickford, D. & Carrasco, L. R. (2015), ‘Quantifying the role of online news in linking conservation research to facebook and twitter’, Conservation Biology 29(3), 825–833.

Proulx, R., Massicotte, P. & PÉpino, M. (2014), ‘Googling trends in conservation biology’, Conservation Biology 28(1), 44–51.

R Core Team (2014), R: A Language and Environment for Statistical Computing, R Foundation for Statistical Computing, Vienna, Austria.

Soriano-Redondo, A., Bearhop, S., Lock, L., Votier, S. C. & Hilton, G. M. (2017), ‘Internet-based monitoring of public perception of conservation’, Biological conservation 206, 304–309.

Statcounter (2017), ‘Search engine market share in china.’, available at: http://gs.statcounter.com/search-engine-market-share/all/china.

