## Supplementary material for "The impact of the “World’s 25 Most Endangered Primates” list on scientific publications and media": Online Appendix (from another version of this preprint: version 1, 2019-02-08)

Table A1. List of species included in the Top25 most endangered primate list from the list of 2000-2002 to 2010-2012 and used in the scientific publication analysis. There is the year of their first mention in the Top25 list and the consecutive mentions in the following Top25 lists. Species names are the current species names (based on IUCN) and not the name used at the time of the Top25 list release.

| Species | First mention | Second mention | Third mention | Fourth mention | Fifth mention | Sixth mention |
|---|---|---|---|---|---|---|
| Ateles fusciceps | 2006 | | | | | |
| Ateles hybridus | 2006 | 2008 | 2010 | | | |
| Ateles hybridus brunneus | 2004 | | | | | |
| Brachyteles hypoxanthus | 2000 | 2002 | 2004 | | | |
| Callicebus barbarabrownae | 2010 | | | | | |
| Cebus flavius | 2010 | | | | | |
| Cebus xanthosternos | 2000 | 2002 | 2004 | | | |
| Cercocebus atys lunulatus | 2000 | 2002 | 2004 | | | |
| Cercocebus galeritus galeritus | 2002 | | | | | |
| Cercocebus sanjei | 2000 | 2002 | 2004 | | | |
| Cercopithecus roloway | 2002 | 2006 | 2008 | 2010 | | |
| Cercopithecus sclateri | 2000 | | | | | |
| Eulemur cinereiceps | 2004 | 2006 | 2008 | | | |
| Eulemur flavifrons | 2008 | 2010 | | | | |
| Galagoides rondoensis | 2006 | 2008 | 2010 | | | |
| Gorilla beringei graueri | 2010 | | | | | |
| Gorilla beringei beringei | 2000 | 2002 | 2004 | | | |
| Gorilla gorilla diehli | 2000 | 2002 | 2004 | 2006 | 2008 | |
| Hapalemur aureus | 2000 | | | | | |
| Hapalemur griseus alaotrensis | 2000 | | | | | |
| Hoolock hoolock | 2006 | 2008 | | | | |
| Hylobates moloch | 2000 | | | | | |
| Lagothrix flavicauda | 2000 | 2006 | 2008 | 2010 | | |
| Leontopithecus caissara | 2000 | 2002 | 2004 | | | |
| Leontopithecus chrysopygus | 2000 | | | | | |
| Leontopithecus rosalia | 2000 | | | | | |
| Lepilemur sahamalazensis | 2006 | | | | | |
| Lepilemur septentrionalis | 2008 | 2010 | | | | |
| Loris tardigradus nycticeboides | 2004 | 2006 | | | | |
| Macaca silenus | 2010 | | | | | |
| Mandrillus leucophaeus | 2000 | | | | | |
| Nomascus hainanus | 2000 | 2004 | 2006 | | | |
| Nomascus nasutus | 2002 | 2008 | 2010 | | | |

| | | | | | | |
|---|---|---|---|---|---|---|
| Nycticebus javanicus | 2008 | 2010 | | | | |
| Piliocolobus pennantii | 2004 | 2006 | 2010 | | | |
| Pongo abelii | 2000 | 2002 | 2004 | 2006 | 2008 | |
| Pongo pygmaeus pygmaeus | 2010 | | | | | |
| Presbytis hosei canicrus | 2004 | | | | | |
| Presbytis natunae | 2002 | | | | | |
| Procolobus badius waldroni | 2000 | 2002 | 2006 | | | |
| Procolobus rufomitratus | 2002 | 2004 | 2006 | 2008 | | |
| Prolemus simus | 2002 | 2004 | 2006 | 2008 | 2010 | |
| Propithecus candidus | 2000 | 2002 | 2004 | 2006 | 2008 | 2010 |
| Propithecus perrieri | 2000 | 2002 | 2004 | | | |
| Propithecus tattersalli | 2000 | | | | | |
| Pygathrix cinerea | 2000 | 2002 | 2004 | 2006 | 2008 | 2010 |
| Rhinopithecus avunculus | 2000 | 2002 | 2004 | 2006 | 2008 | 2010 |
| Rhinopithecus bieti | 2002 | | | | | |
| Rhinopithecus brelichi | 2002 | | | | | |
| Rungwecebus kipunji | 2006 | 2008 | | | | |
| Saguinus oedipus | 2008 | | | | | |
| Semnopithecus vetulus nestor | 2004 | 2006 | 2008 | 2010 | | |
| Simias concolor | 2002 | 2004 | 2006 | 2008 | 2010 | |
| Tarsius tumpara | 2006 | 2008 | 2010 | | | |
| Trachypithecus delacouri | 2000 | 2002 | 2004 | 2006 | 2008 | 2010 |
| Trachypithecus leucocephalus | 2002 | | | | | |
| Trachypithecus poliocephalus | 2000 | 2002 | 2004 | 2006 | 2008 | 2010 |
| Varecia variegata | 2010 | | | | | |

Table A2. List of control species used to account for the overall increase of publication through time for the scientific publication analysis. This species were never included in the Top25 most endangered primate list from the list of 2000-2002 to 2010-2012 and were evenly distributed between the 4 geographical regions (Africa, Asia, Madagascar and Neotropics).

| |
|---|
| Alouatta arctoidea |
| Alouatta palliata palliata |
| Aotus griseimembra |
| Ateles fusciceps rufiventris |
| Ateles geoffroyi ornatus |
| Ateles marginatus |
| Avahi meridionalis |
| Cacajao calvus |
| Cacajao calvus novaesi |
| Callicebus aureipalatii |
| Cebus albifrons albifrons |
| Cebus olivaceus nigrivittatus |
| Cercopithecus ascanius |
| Cercopithecus erythrotis |
| Cercopithecus hamlyni |
| Cercopithecus mitis albogularis |
| Cercopithecus mitis erythrarchus |
| Cercopithecus mitis stuhlmanni |
| Cercopithecus petaurista buettikoferi |
| Cercopithecus petaurista petaurista |
| Cheirogaleus minusculus |
| Chiropotes albinasus |
| Chiropotes chiropotes |
| Colobus guereza guereza |
| Colobus guereza occidentalis |
| Eulemur collaris |
| Eulemur rufifrons |
| Galagoides demidovii |
| Galagoides orinus |
| Galagoides thomasi |
| Hylobates lar |
| Hylobates lar carpenteri |
| Hylobates lar yunnanensis |
| Lepilemur dorsalis |
| Lepilemur grewcockorum |
| Lepilemur hubbardorum |
| Lepilemur leucopus |
| Lepilemur ruficaudatus |
| Macaca leonina |

| |
|---|
| Macaca nemestrina |
| Macaca nigrescens |
| Mandrillus leucophaeus poensis |
| Microcebus marohita |
| Miopithecus talapoin |
| Nasalis larvatus orientalis |
| Nomascus concolor lu |
| Papio ursinus |
| Perodicticus potto |
| Perodicticus potto edwardsi |
| Pithecia pithecia pithecia |
| Pongo pygmaeus morio |
| Presbytis chrysomelas chrysomelas |
| Presbytis frontata |
| Procolobus pennantii bouvieri |
| Pygathrix nigripes |
| Rhinopithecus strykeri |
| Saguinus fuscicollis |
| Saguinus fuscicollis fuscicollis |
| Saguinus fuscicollis illigeri |
| Saguinus fuscicollis nigrifrons |
| Saguinus leucopus |
| Saguinus mystax |
| Saguinus nigricollis graellsi |
| Saimiri boliviensis peruviensis |
| Saimiri sciureus cassiquiarensis |
| Saimiri ustus |
| Sciurocheirus alleni cameronensis |
| Semnopithecus schistaceus |
| Tarsius bancanus natunensis |
| Trachypithecus laotum |
| Trachypithecus obscurus |
| Trachypithecus phayrei |
| Trachypithecus vetulus philbricki |
| Varecia variegata variegata |

Table A3. List of keywords and the scientific and common names of the primate species included in the world's 25 most endangered primates of 2014-2016 officially used for the media penetration analysis. This term expressions were searched in Google News, Twitter, Google Blog Search and Google Trends a month before and a month after its official launch on the 24th November 2015.

| |
|---|
| **top 25 primates** |
| **primates in peril** |
| **endangered primates** |
| **threatened primates** |
| **IUCN primates** |
| Cheirogaleus lavasoensis |
| Lavasoa Dwarf Lemur |
| Hapalemur alaotrensis |
| Lac Alaotra Bamboo Lemur |
| Alaotra Reed Lemur |
| Alaotran Gentle Lemur |
| Lake Alaotra Gentle Lemur |
| Varecia rubra |
| Red Ruffed Lemur |
| Red-ruffed Lemur |
| Lepilemur septentrionalis |
| Northern Sportive Lemur |
| Sahafary Sportive Lemur |
| Propithecus perrieri |
| Perrier's Sifaka |
| Galagoides rondoensis |
| Rondo Dwarf Galago |
| Rondo Bushbaby |
| Cercopithecus roloway |
| Roloway Monkey |
| Procolobus preussi |
| Preuss's Red Colobus Monkey |
| Preuss's Red Colobus |
| Procolobus rufomitratus |
| Tana River Red Colobus |
| Eastern Red Colobus |
| Gorilla beringei graueri |
| Grauer's Gorilla |
| Eastern lowland gorilla |
| Carlito syrichta |
| Philippine Tarsier |
| Nycticebus javanicus |
| Javan Slow Loris |
| Simias concolor |
| Pig-tailed snub-nose langur |
| Pig-tailed Langur |

| |
|---|
| Pig-tailed Snub-nosed Monkey |
| Trachypithecus poliocephalus |
| Golden-headed Langur |
| Cat Ba Langur |
| White-headed Langur |
| Trachypithecus delacouri |
| Delacour's Langur |
| White-rumped Black Leaf Monkey |
| Rhinopithecus avunculus |
| Tonkin Snub-nosed Monkey |
| Semnopithecus ajax |
| Chamba Sacred Langur |
| Kashmir Gray Langur |
| Himalayan Grey Langur |
| Dark-eyed Himalayan Langur |
| Western Himalayan Langur |
| Semnopithecus vetulus nestor |
| Western Purple-faced Langur |
| north lowland wetzone purple-faced langur |
| Nomascus hainanus |
| Hainan Gibbon |
| Hainan Crested Gibbon |
| Hainan Black Gibbon |
| Hainan Black Crested Gibbon |
| Pongo abelii |
| Sumatran Orangutan |
| Cebus kaapori |
| Ka’apor Capuchin |
| Kaapori Capuchin |
| Callicebus oenanthe |
| San Martin Titi Monkey |
| Rio Mayo Titi Monkey |
| Rio Mayo Titi |
| Andean Titi Monkey |
| Alouatta guariba guariba |
| Northern Brown Howler |
| Ateles hybridus |
| Brown Spider Monkey |
| Variegated Spider Monkey |
| Ateles fusciceps fusciceps |
| Ecuadorian Brown-headed Spider Monkey |
| Brown-headed Spider Monkey |

Figure A1. Box plot of effect size of the Causal Impact analysis on the counts of the number of scientific publications containing the species Latin names for each World's 25 Most Endangered Primates' species included in several Top 25 list (multiple mention in red, n= 16) or only once (single mention in green, n= 21) from the 2000-2002 to the 2010-2012 World's 25 Most Endangered Primates' species lists.

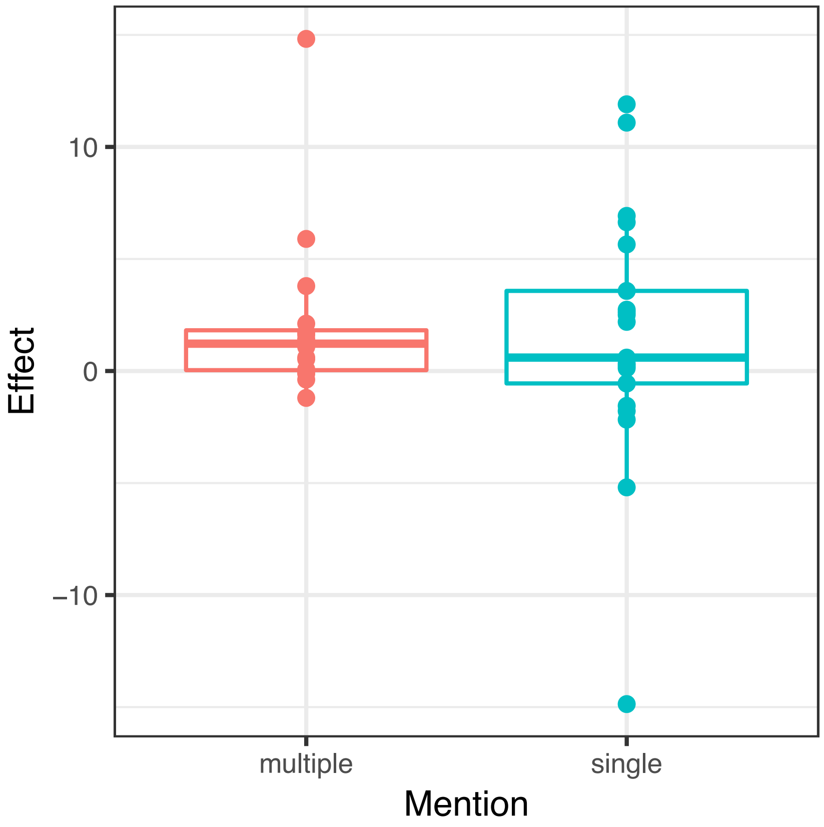

Figure A2. Posterior effect size of Causal Impact analysis for each 37 primate species included in the 6 World's 25 Most Endangered Primates' lists from 2000-2002 to 2010-2012 on scientific publications containing at least once their Latin names. We have used for this figure as control species only species defined as "threatened" (n= 37 species) to account for a general increase in publication through the years on species that are vulnerable to endangerment. Effect size containing only positive values are in blue, containing both positive and negative value are in grey and containing only negative value are in red.

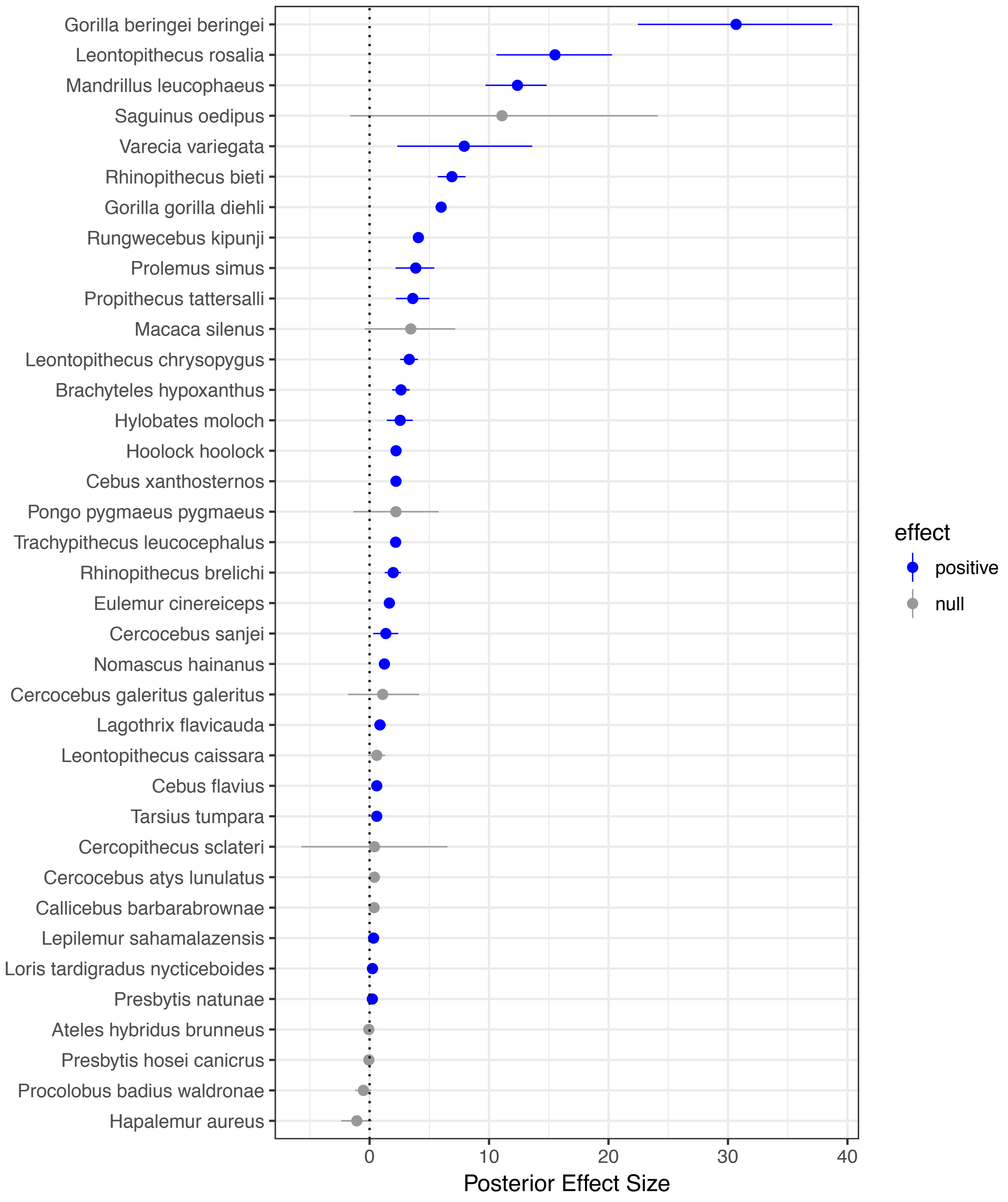
